# Adjunct phage treatment enhances the effectiveness of low antibiotic concentration against *Staphylococcus aureus* biofilms *in vitro*

**DOI:** 10.1101/358879

**Authors:** James Dickey, Véronique Perrot

## Abstract

Phage therapy is drawing more interest as antibiotic resistance becomes an ever more serious threat to public health. Bacterial biofilms represent a major obstacle in the fight against bacterial infections as they are inherently refractory to many types of antibiotics. Treating biofilms with phage has shown promise in a handful of experimental and case studies. However, quantification of the effect of phage combined with antibiotics is needed to pave the way for larger clinical trials. Here we explore the effect of using phage in combination with a total of nine antibiotics, applied simultaneously or as a pretreatment before antibiotics are applied to in vitro biofilms of *Staphylococcus aureus*. Most antibiotics alone were ineffective at low concentration (2×MIC), but the addition of phage to treatment regimens led to substantial improvements in efficacy. At high concentration (10×MIC), antibiotics alone were effective, and in most cases the addition of phage to treatment regimens did not improve efficacy. Using phage with rifampin was also very effective at reducing the outgrowth of resistant strains during the course of treatment.

## Introduction

Bacteriophages (phages), viruses that kill bacteria, have been used to treat bacterial infections since shortly after their discovery just over a century ago (1-3). While research into and use of phage as antimicrobial treatment persisted in a handful of countries, most notably in the Republic of Georgia, the widespread use of antibiotics completely displaced phage therapy in the rest of the world. In recent years, the rise of antibiotic resistance and the spread of pathogen strains resistant to most or all antibiotics (4, 5) has spurred a renewed interest in the use of phages as therapeutic agents (6-10). There is a growing body of evidence demonstrating the effectiveness of phage for treating natural (11) as well as experimental infections in animal models (e.g.,(12-19)). Phage have also been used through the FDA Expanded Access, aka compassionate use, for critically ill human patients infected with bacteria that fail to respond to antibiotics, sometimes with mixed success (20) but occasionally, and quite recently, with impressive outcomes (21-23).

The successful use of phage to treat infections in animals and for compassionate use therapy in human patients is encouraging, and it justifies developing protocols for phage therapy in humans. As is the case for new chemotherapeutic agents, properly controlled clinical trials demonstrating safety and efficacy in human patients are essential. However, clinical trials that implement solely phage for the treatment of infections that are otherwise amenable to antibiotic therapy will face difficult regulatory hurdles (24-26). Alternatively, one could conduct clinical trials to compare treatments combining antibiotics with phage with treatment with antibiotics alone. While there is little or no reason to anticipate adverse reactions to phage in systemic treatment (27, 28), one could start with trials for topical, rather than systemic, treatments of infections and target infections of the skin and mucosa, such as burn wounds, diabetic ulcers, and sinusitis.

The advantage of treating topical infections with the superficial administration of phage and antibiotics over antibiotics alone extends beyond simple regulatory circumstances. Even when they are genetically sensitive to antibiotics, bacteria infecting skin and surfaces of tissues commonly form a biofilm, a polysaccharide matrix that can allow bacteria to be phenotypically refractory to antibiotics (29-32). There is evidence that phage may break open biofilms, making the bacteria within more susceptible to antibiotics (33, 34). Furthermore, there is *in vitro* experimental evidence that combinations of phage and antibiotics are more effective than antibiotics alone for killing biofilm populations of *Pseudomonas aeruginosa*, (35, 36) and *Staphylococcus aureus* (37). Finally, antibiotics and phage may be evolutionarily synergistic. When used in combination, phage may prevent the ascent of antibiotic resistant minority populations, and antibiotics may conversely prevent the ascent of phage resistant bacteria (35, 36), although resistance to phage does not seem to appear readily in *S. aureus*.

In this study, we expand upon the earlier work by Chaudhry et al (36) with *P. aeruginosa* by examining the joint action of antibiotics and phage for treating in vitro biofilms of the Gram positive and common skin pathogen *Staphylococcus aureus*. In addition to using this very different bacterium, we examine a broader range of antibiotics, including but not limited to drugs that are traditionally considered bacteriostatic. We implement these antibiotics against *S. aureus* biofilms at high and low concentrations as a sole treatment, applied simultaneously with phage, and used in sequence following phage treatment. We also explore the effectiveness of phage treatment to suppress the ascent of resistance to rifampin, an antibiotic to which resistance arises quite rapidly, in *S. aureus* biofilms.

The results of this study provide additional support for the potential of the combined use of phage and antibiotics for the treatment of topical infections. They suggest that when applied with phage, low concentrations of antibiotics can be as effective as higher concentrations of antibiotic applied alone, and that phage can prevent treatment failure due to the ascent of antibiotic resistance.

## Materials and Methods

### Bacteria and phage strains

All experiments used the bacterium *S. aureus* Newman given to the lab by William M. Schafer. A single phage isolated from the commercially available Eliava PYO phage cocktail was used. The PYO phage cocktail is routinely use to treat various skin and wound infections In the Republic of Georgia, and contains phages targeted to *Staphylococcus, Streptococcus, Pseudomonas, Proteus* and *E. coli* (6). The single phage isolated from the cocktail is hereafter referred to as PYO phage.

### Phage imaging and sequencing

The phage was imaged by the Robert P. Apkarian Integrated Electron Microscopy Core at Emory University (Figure 1). The left panel shows an intact virion, and the right panel shows a virion with its tail contracted, indicating that the phage we used is likely a member of the family *Myoviridae*. Phage DNA was extracted using the Phage DNA Isolation Kit #46500 from Norgen Biotek Corp., and sequenced by the Molecular Evolution Core of the Parker H. Petit Institute for Bioengineering and Bioscience at the Georgia Institute of Technology. The sequence was deposited in NCBU under the BioProject accession number PRJNA477834. It clusters with Staphylococcus virus G1 (accession number AY954969) published in (38).

**Figure 1:**
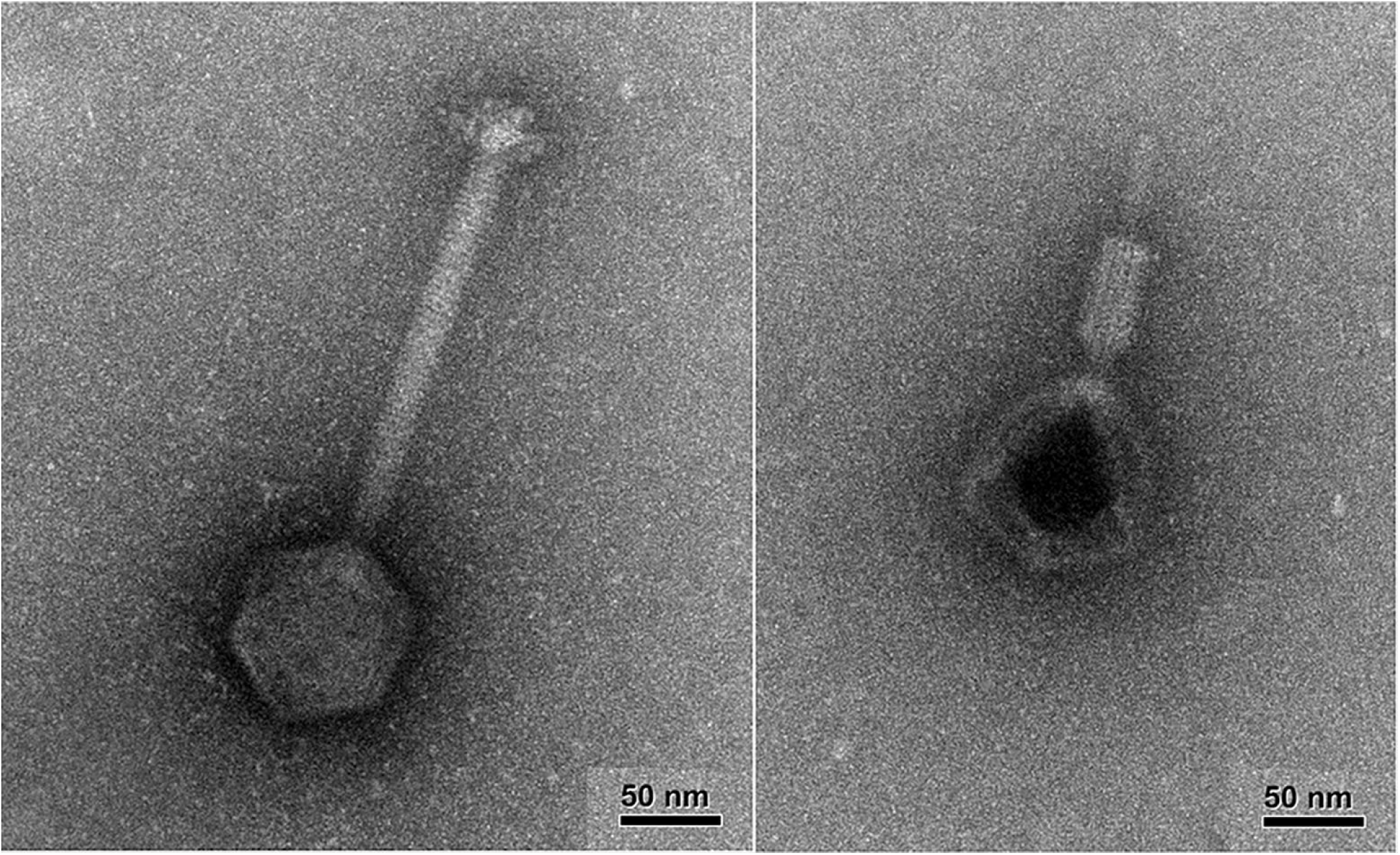
Transmission Electron Microscopy images of the PYO phage. The images were taken on a JEOL JEM-1400 120kV TEM. Left: intact virion. Right: virion with its tail contracted.

### Culture and sampling media S. aureus

Newman was grown in Muller Hinton II (MHII) broth (Difco). Bacterial densities were determined by serial dilution in saline and plating on Lysogeny Broth (LB) hard (1.6%) agar. Phage densities were estimated by mixing the serially diluted culture with a lawn of *S. aureus* Newman (0.1mL of a 1 in 10 dilution of a fresh overnight culture), adding 1.5 mL of soft (0.65%) LB agar, and pouring the mixture on the surface of 1% agar LB plates. When added to biofilms,p hage was added at ∼4e6 pfu/mL, at an MOI of ∼0.1.

### Antibiotics and MIC estimation

Antibiotics were purchased from Sigma (gentamicin (GEN), oxacillin (OXA), vancomycin (VAN), and tetracycline (TET)), AppliChem (ciprofloxacin (CIP), rifampin (RIF)), TCI (daptomycin (DAP)), MP Biochemicals (erythromycin (ERM), and Chem-Impex International (linezolid (LIN)). The minimum inhibitory concentration (MIC) for each drug was determined by the standard two fold dilution protocol (39). In the expriments, each antibiotic was used at 2×MIC and 10×MIC. We chose these antibiotics to have a broad range of pharmacodynamic properties. The first eight antibiotics listed above are grouped by their pharmacodynamic characteristics: in classic assays of killing dynamics in liquid cultures at super MIC concentrations, CIP, DAP and GEN kill rapidly (fast-acting bactericidal antibiotics); OXA and VAN kill more slowly (slow-acting antibiotics); and LIN, ERM and TET prevent growth (bacteriostatic antibiotics) (40).

### Biofilm establishment and preparation, treatment, and sampling

To establish biofilms of *S. aureus*, 24-well tissue culture plates (TPP; well diameter: 15 mm, surface of the biofilm: 177 mm^2^) were inoculated with 2 mL MHII broth with ∼ 10^6^ 121 cfu/mL of an overnight culture of *S. aureus* Newman. Plates were incubated at 37°C without shaking for 72 h to establish a biofilm on the bottom of the wells. After 72 h incubation, each well was prepared by removing the media using an aspirator and washed twice with 2 mL saline. After removal of the last saline wash, 1 mL of broth containing antibiotic (antibiotic-only controls), phage (phage-only controls and first phase of sequential treatments), or both (simultaneous treatments (SIM)) or neither (untreated controls), was added to each well and incubated without shaking for 48 h. In the case of sequential treatment (phage first, antibiotic later (SEQ)),p hage was added, plates incubated without shaking at 37°C for 24 h, at which time the appropriate amount of antibiotic (2x or 10x MIC) was added to each well in saline, and incubated without shaking at 37°C for an additional 24 h for a total treatment time of 48 h.

To estimate the density of bacteria and phage after the 48 h incubation, the biofilms at the bottom of the wells were disrupted with the flat end of a sterile wooden stick; the walls were not disrupted. The bacteria and phage in the wells were further mixed by taking up and returning 100 μL with the pipette. The densities of bacteria and phage in these suspended cultures were then estimated by serial dilution and plating. When desired, the density of bacteria resistant to RIF was estimated by plating on plates containing 10×MIC of RIF. When most of the bacteria were able to form colonies on RIF-containing plates the population was classified as resistant; otherwise it was classified as sensitive.

The initial, pre-treatment densities of bacterial in the biofilms were estimated as above by adding 1 mL of saline, rather than broth, to the 72 h prepared biofilms. Since the sampling of a biofilm requires its disruption, each estimate of densities was taken from a different well. In addition, the density of phage added to the well was estimated by plating.

### Experimental design and sample size

For all antibiotics except RIF, each combination of a given antibiotic at a given concentration, used alone or with phage (either simultaneously or sequentially) was run three separate times with three replicate wells each time. Because of various mishaps, the final number of data points for each combination was between 6 and 9. For all antibiotics except RIF, the antibiotic free controls (no treatment and phage-only controls) were pooled, with a total of between 16 and 30 wells each. Thus, the results presented in Figures 2 and 3 all have the same antibiotic free controls. Experiments with RIF were done at a later time, and were run two separate times with 10 replicates each time for each combination of RIF at 2×or 10×MIC, used alone or with phage (either simultaneously or sequentially). Additionally, they had their own untreated and phage only controls (a total of 6 wells for each control).

**Figure 2:**
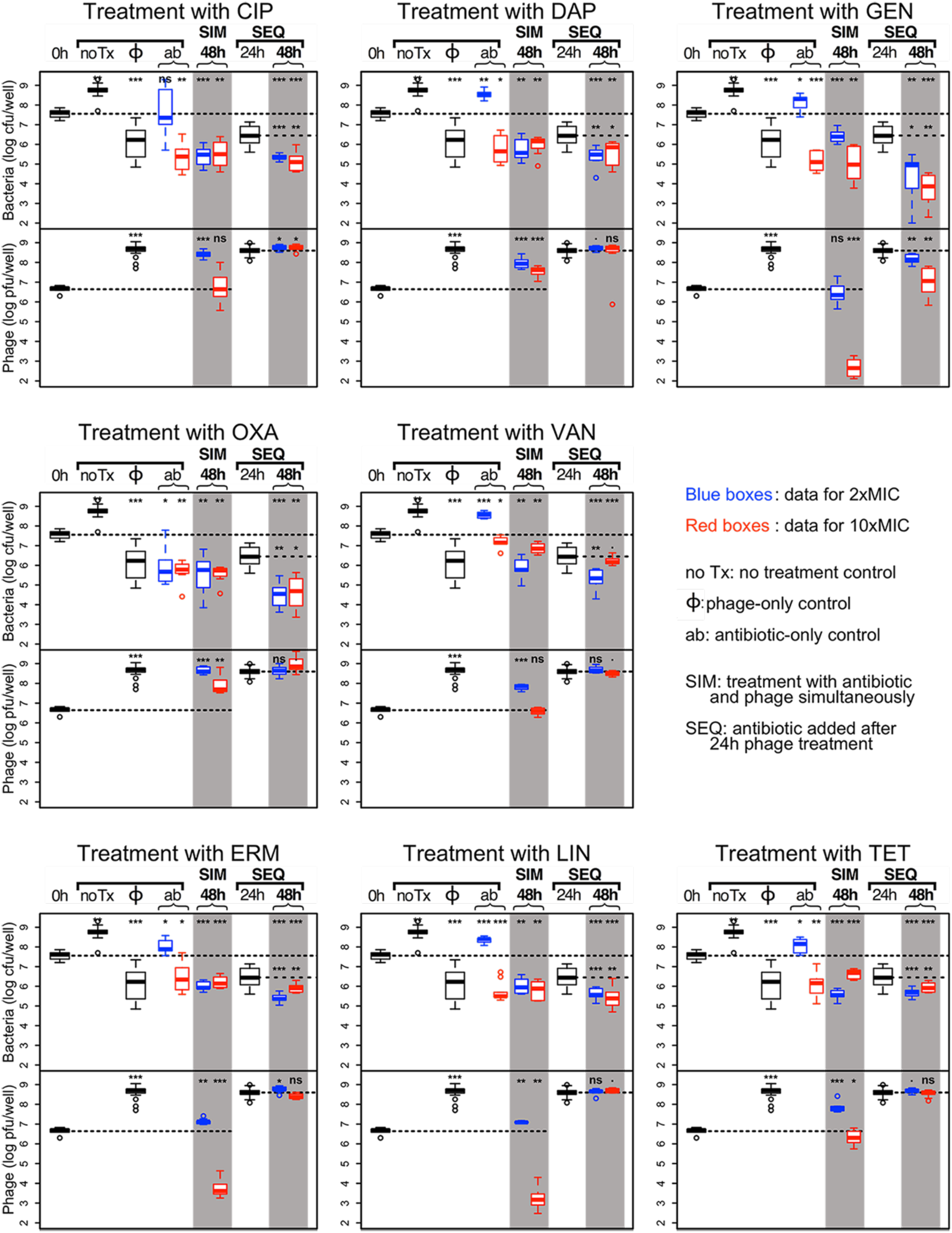
Combined action of 8 antibiotics and phage. For each plot, top panel: bacterial densities; bottom panel: phage densities. Blue boxes: treatment with 2×MIC of the antibiotic; red boxes: treatment with 10×MIC of the antibiotic. Densities after 48h of combined antibiotic and phage treatments are shown in the shaded columns. For each plot we use the same controls (the left side of the plot): (i) untreated, or treated with phage only (same data for all 8 antibiotics), and (ii) treated only with the noted antibiotic for that plot. For each plot, the results for the simultaneous treatment (SIM) with phage and 2× and 10×MIC of the antibiotic are shown in the middle shaded column. The sequential treatment (SEQ), where the cultures were first treated with phage for 24 h then treated with the antibiotic for another 24 h, the bacterial and phage densities at 48 h are shown on the right shaded column. The bacterial and phage densities after 24 h exposure to phage are shown between the two shaded columns. The dashed lines across the panels indicate the initial densities of bacteria and phage. The short, horizontal broken lines in the 3 columns to the right for the sequential treatment denote the densities of bacteria (top panel) and phage (bottom panel) after 24 h exposure to phage. The significance levels of the two-tailed t-tests comparing the densities at 48 h with initial densities and with densities at the time antibiotic was added in SEQ treatment are as follows: ***: p<0.00001, **: p<0.001, *: p<0.05; ·: p<0.1 and NS: p>0.1.

**Figure 3.**
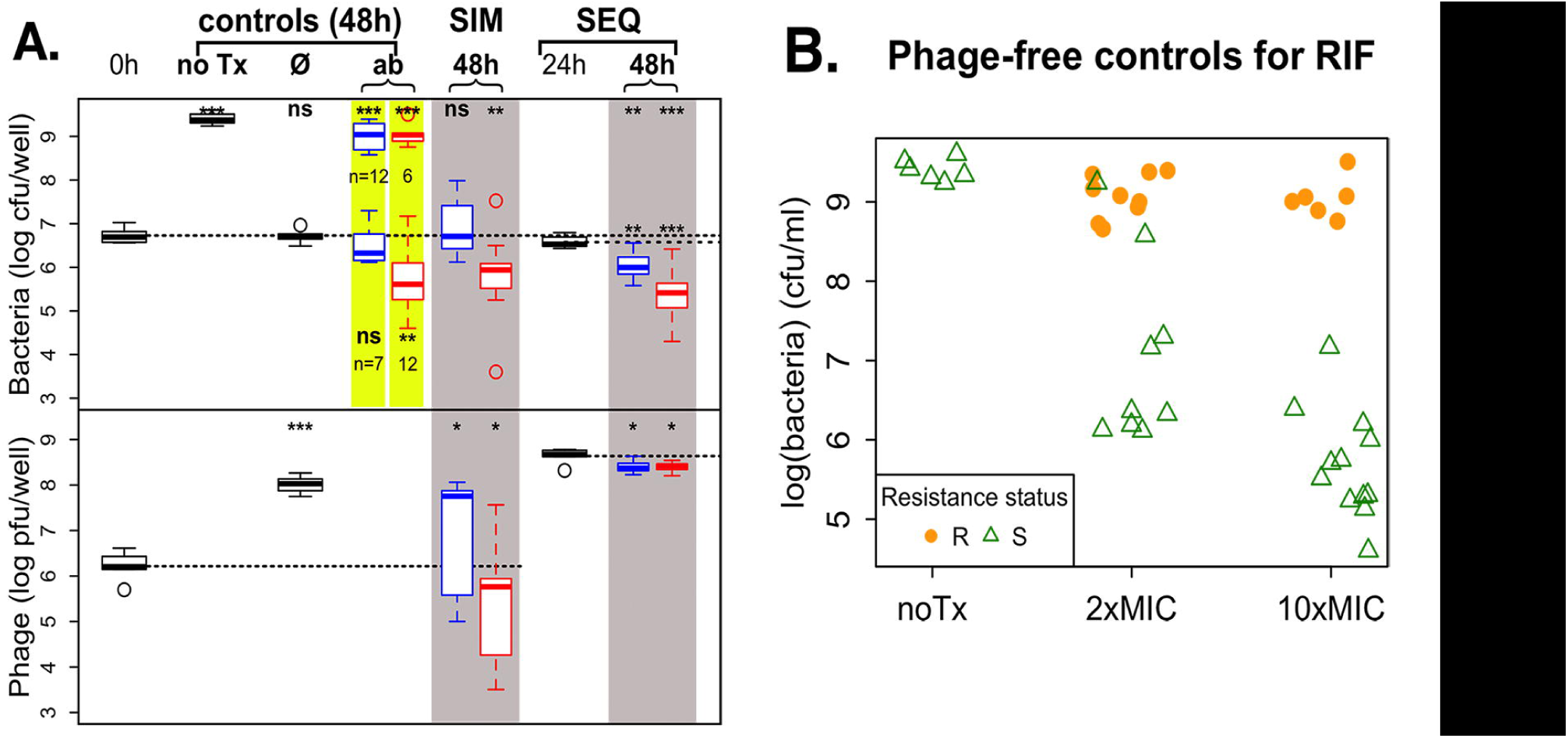
Treatment of *S. aureus* biofilm with RIF and phage. **A.** Combined action of phage and RIF on *S. aureus* biofilms. The legend is the same as for Figures 2 and 3. 20 wells were used for antibiotic-only controls, and simultaneous and sequential treatments, at each antibiotic concentration. The 2× and 10×MIC RIF-only control data are highlighted in yellow: some wells became turbid over the 48h of incubation (bacterial density ∼ 1e9; data is the same as the points at the top of figure 4B), and some wells remained clear (bacterial density ∼ 1e5 - 1e7; data is the same as the points at the bottom of figure 4B). The number of wells in each category is indicated below the corresponding box. **B.** Resistance status of phage-free controls. Bacterial densities are shown for each well, in solid orange dots for resistant wells and open green triangles for sensitive wells. Note that some of the cultures dosed with 2×MIC of RIF reached high densities (>1e8 cfu/mL) even though they appear to be sensitive to RIF when tested on plates containing 10×MIC RIF.

### Measure of the efficacy of treatment and statistical analyses

For our experiments, we define an effective treatment as one that leads to a decrease in the density of bacteria recovered from a well relative to the density before treatment (initial density). In our protocol, we do not remove the treatment media before disrupting the biofilm. Instead, the biofilm is mixed with the planktonic population for sampling at the end of treatment. A net decrease in bacterial density of the culture after time 0 indicates that the treatment was effective at reducing the bacterial density in the biofilm. In addition, a decrease in culture density would also indicate that treatment prevented the outgrowth of a planktonic cells. As could be expected, there is a 1 to 2 log increase in bacterial density when only broth is added to the biofilm (untreated controls; see Figures 2 and 3, and Tables 1 and 2). Thus, if there is no significant change in density after 48 h, the treatment does not reduce the bacterial density, but it does prevent growth of the biofilm or the outgrowth of a planktonic cells, or some combination of both.

**Table 1:**
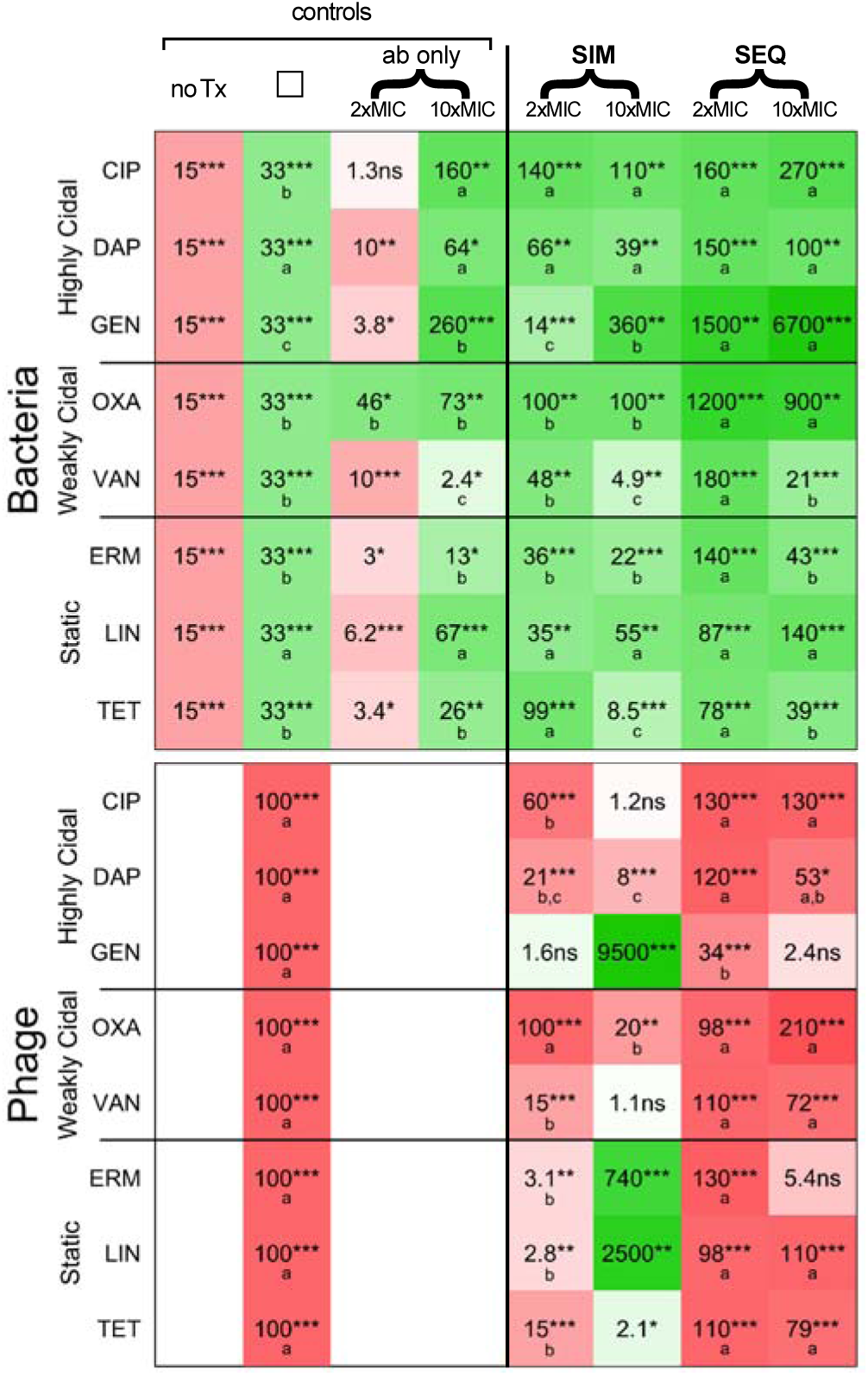
Summary of the effects of different treatments on the densities of *S. aureus* and phage. This table shows the changes in bacterial density (top panel) and phage density (bottom panel) over 48 h compared with initial densities for 8 antibiotics and the phage and antibiotic treatments on the right, and for the various controls on the left (no Tx=untreated; ø= phage only; ab only=antibiotic only). Cells in green hues show a decrease compared with initial densities; cells in red hues show an increase; the intensity of the colors indicates the magnitude of the change in density (the more intense the color, the greater the change). The number in each cell indicates the magnitude of the change, and the stars indicate the level of significance of the t-test to determine if the change was significant, as follows: ***: p<0.00001; **: p<0.001; *: p<0.05; ·: p<0.1; NS: p≥0.1. In the top panel, green cells indicate that the treatment was effective. Considering only effective treatments, the letters in the green cells indicate the groups of significance within each line: for a given antibiotic, values with the same letter are not significantly different from each other, but values with different letters are. In the bottom panel, red cells indicate that the phage increased in density. Considering only treatments that led to a significant increase in phage, letters in the red cells indicate the groups of significance within each line: for a given antibiotic, values with the same letter are not significantly different from each other, but values with different letters are.

**Table 2:** Summary of the effects of different treatments on the densities of S. aureus and phage using RIF. The top row of the table shows the changes in bacterial density over 48 h compared with initial densities. For the RIF-only controls, the values for the wells that did not become turbid are shown in the top row, and the values for the wells that became turbid are shown on the second row. Otherwise, same legend as Table 1.

|  | controls |  |  |  | SIM |  | SEQ |  |  |
| --- | --- | --- | --- | --- | --- | --- | --- | --- | --- |
| | noTx | $\phi$ | ab only | | 2xMIC | 10xMIC | 2xMIC | 10xMIC | |
| **Bacteria** | 470*** | 1ns | 1.6ns | 11**<br>a,b | 1.5ns | 8.3**<br>a,b | 4.6**<br>b | 23***<br>a |  |
|  |  |  | 190*** | 210*** |  |  |  |  |  |
| **Phage** |  | 62***<br>a |  |  | 5.3*<br>b | 6.3* | 150***<br>a | 150***<br>a |  |

To determine the statistical significance of the change in bacterial or phage density after 48 h treatment, we performed 2-tailed t-tests comparing the initial density and the density after treatment. For each antibiotic, we wanted to determine if some treatments were more effective than others. To do this, we ran an analysis of variance including only the treatments that were effective (i.e., treatments that led to a significant decrease in bacterial density compared with initial densities). Similarly, we wanted to determine if some treatments led to greater phage growth, so we ran an analysis of variance including only the treatments that lead to significant phage growth.

All statistical analyses were performed on log-transformed bacterial or phage densities. All analyses were done using R in its RStudio incarnation (41). In Figures 2, 3, and 4, the results are represented by standard box-and-whisker plots, where in each box the black line shows the median density of bacteria and phage, the box illustrates the range for 50% of the data (IQR), the whiskers show 99.3% of the data, and the open circles show the points that fall outside of the whiskers.

**Figure 4:**
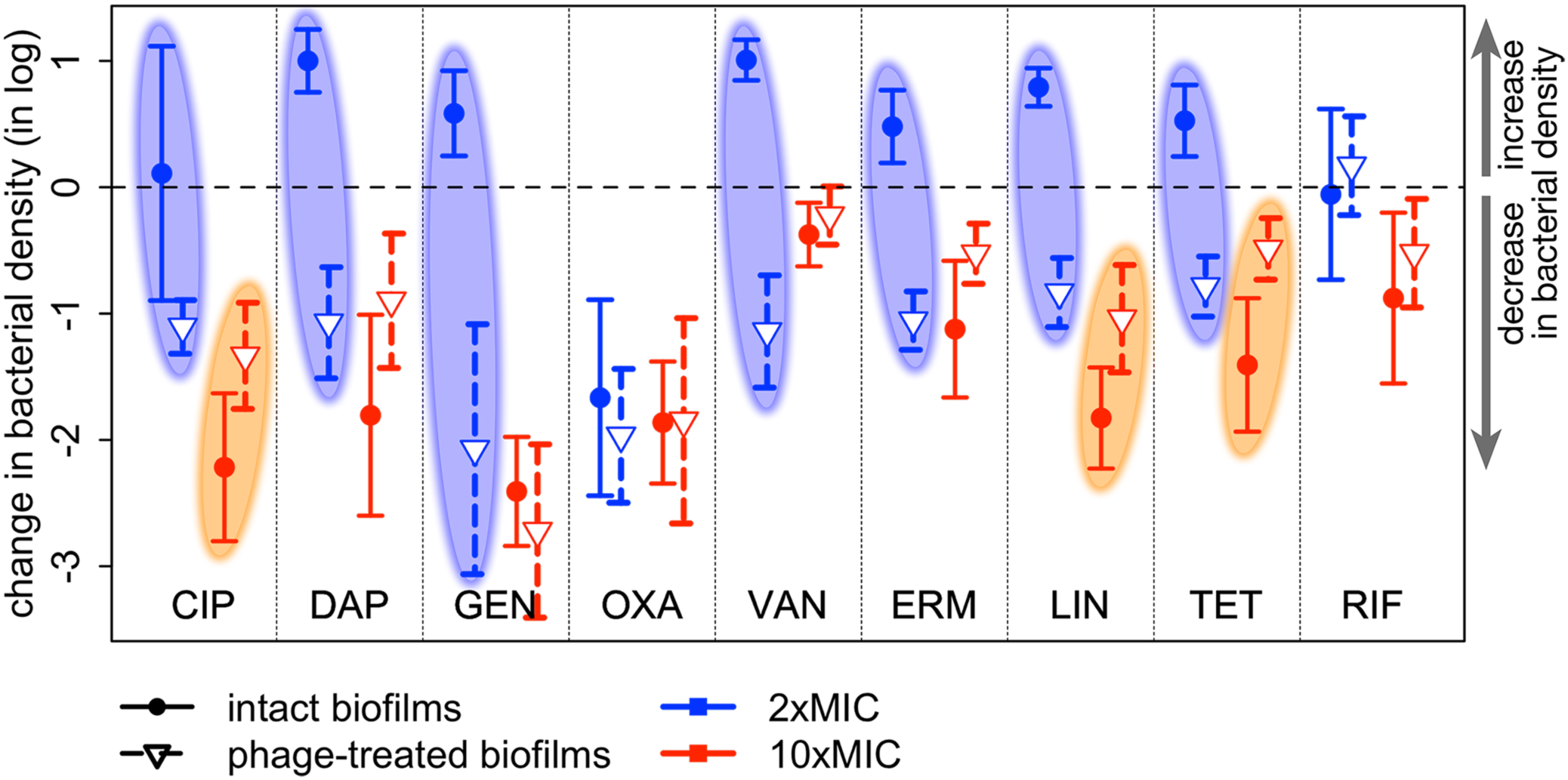
Change in bacterial density after adding antibiotics to intact or phage-treated biofilms. The change in bacterial density (log(final density) – log(initial density)) is shown for all 9 antibiotics used in this study at both antibiotic concentrations. The symbols indicate the difference in means, and the error bars indicate the 95% confidence interval obtained from t-tests. Values above the dashed line show an increase in bacterial density during the course of treatment; values below the dashed line a decrease. Antibiotics added to intact biofilms are shown in solid dots and solid lines (initial densities = bacterial densities in prepared biofilms, final densities = antibiotic-only controls); antibiotics added to biofilms exposed to phage for 24 h are shown in open triangles and dashed lines (initial densities = bacterial densities after 24 h of phage-only treatment, final densities = sequential treatment at 48 h).

## Results

### 1 Bacterial strain characterization

For *S. aureus* Newman, the MICs for each drug were, in μg/mL: CIP, 0.1875; DAP, 2; GEN, 0.75; OXA, 0.1875; VAN, 1.5; ERM, 0.375; LIN, 1.5; TET, 0.1875; RIF, 0.01. Unlike *E. coli* or *P. aeruginosa* and their lytic phages, *S. aureus* does not seem to readily evolve resistance to its phage. We do not observe the appearance of bacteria resistant to the PYO phage in standard culture: resistant colonies do not appear on phage-treated lawns, nor do cultures of *S. aureus* with phage become turbid overnight, as is commonly observed with *E. coli* and *P. aeruginosa* and their phages.

### 2 Combination treatment with phage and antibiotics

In this experiment, we measured the bacterial (and phage, where applicable) densities in prepared biofilms of *S. aureus* before (initial density, shown as 0h on Figure 2) and after 48 hours of treatment with combinations of phage and antibiotics. As controls, we also measured bacterial and phage densities after 48h of treatment with phage, antibiotics, or neither (untreated control). As sampling is destructive, a set of wells were sacrificed to estimate the initial density in biofilms at the time of treatment. The same initial density data is used for all the 8 antibiotics in this experiment. In addition, we use the same data for the untreated and phage-only controls for all 8 antibiotics.

The results of all the antibiotics except RIF are presented in Figure 2 and summarized in Table 1.

For the convenience of the reader we describe in some details one representative plot in Figure 2; we arbitrarily chose DAP for this purpose. In untreated controls, the viable cell density of *S. aureus* increased by 15 fold relative to the initial density. In the phage-only controls, the cell density was reduced 33 fold, and the phage density increased by 100 fold. At 2×MIC, DAP alone was not effective in reducing the viable cell density of bacteria. DAP at 10×MIC and simultaneous and sequential treatments at both antibiotic concentrations were effective in significantly reducing the viable cell density of *S. aureus* in culture, and not significantly different from each other (second line of top panel of Table 1). DAP added at both concentrations after 24 h of phage treatment (SEQ) was able to further reduce the bacterial density: the blue and red boxes in the right shaded column are significantly below the dashed line showing the bacterial density after 24 h of phage exposure (middle plot of the top row of Figure 2). This observation holds even when the antibiotic was added at 2×MIC, a concentration at which DAP alone was unable to prevent the outgrowth of an intact biofilm.

There was a significant increase in phage density in all the combined therapy experiments with DAP (second line of the bottom panel of Table 1). On the other hand, the increase in phage density in simultaneous treatment with DAP was significantly less than in the absence of this antibiotic. With sequential treatment, the density of phage increased to roughly that observed in the absence of this antibiotic.

In the following, we summarize the results of these experiments

#### (1) Untreated and phage-only controls

(i) In absence of phage or antibiotics, the density of bacteria increased by 15 fold.

(ii) Phage reduced the viable cell density in the culture by 33-fold, and the density of phage increased by two orders of magnitude

#### (2) Antibiotic-only controls

(i) In the absence of phage and at 2×MIC, OXA was the only antibiotic that effectively killed *S. aureus*; it reduced the viable cell density in the culture by nearly 50 fold. None of the seven other antibiotics at 2×MIC significantly reduced the viable cell density of *S. aureus* in the culture.

(ii) At 10×MIC, all eight antibiotics significantly reduced the density of *S. aureus* in the culture. This included the officially bacteriostatic antibiotics (ERM, LIN and TET), which are expected to prevent growth rather than kill. CIP and gentamicin were more effective than phage alone, while VAN less effective than phage alone. The other five antibiotics were not significantly more effective than phage alone.

#### (3) Antibiotics and phage — Simultaneous treatment

(i) In combination with 2×MIC of all 8 antibiotics, simultaneous treatment with phage reduced the viable cell density of bacteria. Adding phage did not improve the performance of the already effective OXA. For CIP and TET, the efficacy of the combination (CIP+phage: 140; TET+phage: 99, Table 1) was significantly greater than that of phage acting alone (33). Since these antibiotics alone at 2×MIC were unable to kill bacteria (CIP) or prevent outgrowth (TET), this is a case of synergy, where the efficacy of combination of phage and antibiotic is significantly greater than the sum of the individual efficacies. For the other six antibiotics, the efficacy of phage alone and phage with 2×MIC of the antibiotic are not significantly different.

(ii) There was significant phage growth when antibiotics were added with phage at 2×MIC for all antibiotics except GEN, although the phage increased less in the presence of most antibiotics than in the phage-only control. Only OXA had no effect on phage growth.

(iii) Simultaneous treatment with phage and 10×MIC of all 8 antibiotics was effective at reducing bacterial densities in the cultures, but not significantly more effective than the antibiotic alone at 10×MIC for 7 out of the 8 antibiotics. For TET, adding phage actually reduced the efficacy of the antibiotic. When comparing with phage alone, CIP and GEN at 10×MIC with phage were more effective. There was antagonism between VAN and TET at 10×MIC and the phage: the efficacy of the combination (VAN+ phage: 4.9; TET+ phage: 8.5; Table 1) was significantly smaller than that of the most effective agent (phage acting alone: 33) were less effective. Adding 10×MIC of the remaining four antibiotics to treatment with phage did not increase treatment efficacy.

(iv) When added at 10×MIC simultaneously with phage, most antibiotics either prevented phage growth (CIP, VAN, TET) or led to massive decreases in phage density (GEN, ERM, LIN). Only DAP and OXA allowed some modest phage growth.

#### (4) Antibiotics and phage — Sequential vs. simultaneous treatment

(i) For all antibiotics, treatments with phage first, then antibiotic at either concentration were effective at reducing bacterial density. For VAN, ERM and TET, treatment with 10×MIC was significantly less effective than treatment with 2×MIC, while for the other five antibiotics there was no significant difference between the antibiotic concentrations.

(ii) The antibiotics that were not able to prevent bacterial growth of intact biofilms (antibiotic only controls) at 2×MIC, that is, all antibiotics but OXA, were effective at reducing the bacterial density of cultures treated with phage for 24 h.

(iii) For CIP, DAP and LIN, simultaneous treatment with 2× or 10×MIC of the antibiotic was as effective as sequential treatment at either antibiotic concentration.

(iv) GEN and OXA were more effective when added to phage treated cultures (sequential treatment) than when added at the same time as phage (simultaneous treatment).

(v) VAN and TET showed a similar pattern: when used in the presence of phage in both treatment regimens (SIM and SEQ), using the high antibiotic concentration led to significantly less reduction in bacterial density. For TET, the most effective treatment was with 2×MIC with both treatment regimens. For VAN, the most effective treatment was sequential treatment with 2×MIC. Similarly, ERM was most effective when used in sequential treatment at 2×MIC.

(vi) When antibiotics were added in sequential treatment (after 24 h of phage exposure), the phage had grown by two orders of magnitude. There was generally very little change in phage density after antibiotics were added at either concentration, as can be seen in Figure 2, bottom panel for each plot, right shaded column. The only exception was GEN, where there was a small but significant decrease in phage density when the antibiotic was added at 2×MIC, and a larger decrease when added at 10×MIC.

### 2 Phage preventing the ascent of antibiotic resistance

We ran experiments similar as those described above with the additional goal of monitoring the appearance of resistance to the antibiotic RIF. RIF is a potent killer of *S. aureus* at very low concentrations (MIC=0.01 μg/mL), but it is rarely if ever used in the clinic because resistance to the antibiotic emerges very quickly. In our hands, an overnight culture of *S. aureus* treated with RIF at super-MIC concentrations is turbid by morning and the majority population resistant to RIF, a phenomenon we never observed for any other antibiotics. We modified the design of the experiment to include larger sample sizes (a total of 20 wells for each treatment group). We never observed the appearance of resistance to the phage in all the experiments reported in this paper, so we conclude that resistance to phage is not an issue at the time frame of a few days in the biofilm cultures we used. The results are presented in Figure 3 and summarized in Table 2.

The work with RIF was done separately from that with the other 8 antibiotics, with its own set of antibiotic-free controls. We notice that the initial bacterial density of the prepared biofilms was about 6 times lower than for the first 8 antibiotics despite our using the same experimental protocol. The phage density was about the same, so the MOI was slightly higher for the RIF treatment. In this case, phage alone was able to prevent the net growth of bacterial cultures, but unable to significantly reduce the bacteria below their initial densities. Phage densities increased significantly, but slightly less than they did in the first experiment.

The RIF-only controls at both concentrations presented a pattern we didn’t see in any of the other 8 antibiotics: there were two, rather than one, qualitative outcomes. Some wells became turbid, which is equivalent to treatment failure, with bacterial densities greater than 1e8 cfu/mL, and the other wells remained clear, with bacterial densities smaller than 1e8 cfu/mL. Densities in the turbid wells were very close to the density in the untreated controls. For the wells that remained clear (7 out of 19 wells for 2×MIC and 6 out of 18 wells for 10×MIC), rifampin alone at 2×MIC did not significantly reduce the viable cell density of bacteria, as observed with 7 of the other 8 antibiotics (OXA being the exception). RIF alone at 10×MIC was effective at reducing bacterial density, albeit its efficacy was in the range of the weakly bactericidal and bacteriostatic antibiotics (see Table 1).

We assessed the resistance status of the bacteria in the untreated wells, and wells treated with RIF alone (Figure 4B). Not unexpectedly, the bacteria in the untreated wells were sensitive to RIF, as were the bacteria in wells treated with RIF that remained clear. The dominant populations of bacteria in the wells treated with 10×MIC RIF that became turbid were resistant, as were most of the dominant populations in wells treated with 2×MIC RIF that became turbid. In addition, two of such wells appeared sensitive, possibly because the bacteria were tested on plates containing 10×MIC of the antibiotic. The dominant populations in these two wells may have been resistant to 2×MIC but unable to form colonies in the presence of 10×MIC RIF.

When RIF was used simultaneously with phage, its effect was similar to that of RIF alone, but without treatment failure. RIF at 2×MIC with phage did not lead to net growth or death, and RIF at 10×MIC with phage had the same efficacy as RIF at 10×MIC alone when resistance did not arise. RIF was the only antibiotic out of the 9 we tested where simultaneous treatment with 2×MIC of the antibiotic was not effective at reducing bacterial density.

When RIF was added to phage treated wells (sequential treatment), both concentrations significantly reduced bacterial densities, with 10×MIC being significantly more effective than 2×MIC. As in simultaneous treatment, we did not observe any treatment failure in sequential treatment. Overall, there were 17 treatment failures out of 37 wells treated with RIF only, and none out of 80 wells treated with a combination of RIF and phage.

## Discussion

Bacteria in biofilms are notoriously difficult to kill with antibiotics, and biofilms represent a significant problem in treating many bacterial infections (29-31, 42, 43). One of the most cited factors contributing to this problem is the extracellular matrix that comprises the structure of the biofilm (44, 45). This polysaccharide matrix is thought to reduce the exposure of bacteria in the biofilm to the antibiotic, resulting in bacteria that are exposed to a lower, and thus less effective, concentration of antibiotic.

Phages hold some promise in treating biofilm infections. As naturally occurring bacterial viruses, phages may have evolved mechanisms to infect bacteria living in a biofilm (33, 46). Indeed, direct effect of phage on biofilm structure has been demonstrated in *Klebsiella* (47), *P. aeruginosa* (16, 48) and *S. aureus* (49). Using phage alongside antibiotics may improve the efficacy of conventional antibiotic treatment in topical infections, such as burns, skin ulcers, and sinusitis. However, only a handful of experimental studies have quantified the interactions between phage, antibiotics, and bacteria in biofilms. Chaudhry et al. (36) investigated how a pair of phages improved the antimicrobial effects of five bactericidal antibiotics for the control of biofilms of the Gram-negative bacterium *P. aeruginosa* in vitro. In general, they observed that simultaneous use of phage and antibiotics improved the killing of bacteria for two antibiotics (ceftazidime and colistin), while using antibiotics after phage pre-treatment (sequential treatment) improved killing by another two antibiotics (GEN and tobramycin (TOB)). Kumaran et al. (37) studied the combined effect of five antibiotics, three bactericidal and two bacteriostatic, and one phage on biofilms of *S. aureus*. In general, they find that the greatest reduction in bacterial density in the biofilms is obtained when the biofilms are treated with phage first for 24 h, then with antibiotics for 24 h (sequential treatment). However, Kumaran et al. used very high concentrations of antibiotics, with two-fold increasing concentrations starting at 8×MIC for VAN and LIN, 64×MIC for cefazoline and TET, and 256×MIC for dicloxacillin. This large difference with the antibiotic concentrations we used (2 and 10×MIC) make direct comparisons difficult.

This study initially served to extend the work of Chaudhry et al., as we sought to examine the generality of the effectiveness of combined phage and antibiotic treatment against the common skin pathogen and Gram positive *S. aureus*. As with the previous work, antibiotics were applied at the same time as phage (simultaneous treatment) or after phage pre-treatment (sequential treatment), but with a broader range of nine antibiotics that included some bacteriostatic antibiotics. To allow us to fully gauge the effectiveness of our treatment regimens in eliminating bacteria within a biofilm, we further modified Chaudhry et al.’s protocol by using a different measure of efficacy. In our modified protocol, we estimated the bacterial density in prepared biofilms before treatment, and after 48 h of exposure to antibiotic, phage, both, or neither (untreated controls). Instead of comparing bacterial densities after treatment with densities in untreated controls, we compared them with bacterial densities before treatment. Our measure of treatment efficacy is thus the change in bacterial density during the course of treatment. This more conservative estimate allows us to ascertain the ability of treatment regimens to penetrate and kill bacteria within a biofilm. Samples taken after treatment consisted of both biofilm and planktonic bacteria, whereas the samples taken at the time of treatment consisted of only biofilm bacteria. Any treatment that resulted in lower bacterial densities than were in biofilms before treatment must have been effective at penetrating and killing bacteria within the biofilm. In addition, by estimating the bacterial density after 24 h of phage treatment, we could estimate the efficacy of antibiotics on phage-treated biofilms (see Figure 4 below).

The blue ovals highlight the 7 antibiotics used at 2×MIC that did not reduce the bacterial density when applied to intact biofilms (change in density at or above the dashed line) but significantly killed bacteria when added to phage-treated biofilms (change in density below the dashed line). The orange ovals highlight the 3 antibiotics where pretreatment with phage likely reduced the efficacy of 10×MIC antibiotic.

Our results are consistent with that of Chaudhry et al, except that we observed better killing with antibiotic and phage combinations only for low antibiotic concentrations. Such concentrations would otherwise be ineffective or marginally effective, but the inclusion of phage in these treatments led to substantial killing of bacteria in *S. aureus* biofilms. However, unlike the general findings of Chaudhry et al, increasing the concentration five-fold led to no additional killing of bacteria when used with phage. An exception to this pattern is GEN, where in our experiments, GEN was still qualitatively, albeit not always significantly, more effective at high compared with low concentration within each treatment protocol. In some cases, increasing antibiotic concentration led to a decrease in the efficacy of treatment, such as with VAN and TET in simultaneous treatments, and for VAN and ERM in sequential treatments. This is in line with the observations of TOB against *P. aeruginosa* by Chaudhry et al.

Generally speaking, antibiotics at low concentration (2×MIC) were unable to kill bacteria in biofilms while the same antibiotics were able to kill at high concentration (10×MIC). Interestingly, this observation held for bacteriostatic as well as bactericidal antibiotics. Using antibiotics and phage at the same time (SIM treatments) improved killing over antibiotics alone or phage alone in 2 out of 18 cases (9 antibiotics×2 concentrations). For these two cases (CIP and TET at 2×MIC), the antibiotic alone did not reduce bacterial densities, so we can conservatively put their efficacy at 0, if not as a negative number. Since the efficacy of the combination was significantly greater than that of phage alone, there is synergy between antibiotic and phage. There were two cases of antagonism, VAN and TET at 10×MIC, where the efficacy of the combination was smaller than that of the most effective agent acting alone. For all other cases (14 out of 18), the efficacy of the combination was not significantly different from that of the most effective agent acting alone: in some cases, the phage appeared to do most of the killing, in other cases it was the antibiotic. This would indicate that at least in some situations, combined phage and antibiotic would be generally more effective than antibiotic alone, not so much because of synergy between the two agents (only 2 in 18 cases) but because either agent alone is effective (12 out of 18 cases), and antagonism between the two is infrequent (2 out of 18 cases).

Our experimental design further allowed us to check if phage pre-treatment could alter the susceptibility of biofilm-bound bacteria to antibiotics. To answer this question, we estimated the efficacy of the antibiotics when added to intact biofilms (antibiotics-only controls), and when added to phage-treated biofilms (sequential treatment regimens). These efficacies are summarized in Figure 4. When used at 2×MIC, 7 out of 9 antibiotics showed a greatly increased efficacy against phage-treated biofilms compared with intact biofilms: for all but OXA and RIF, the antibiotic was able to kill effectively phage-treated biofilms, while the same concentration didn’t kill intact biofilms (shown in blue ovals on Figure 4). OXA was able to kill intact biofilms, and RIF was not, but pre-treatment with phage did not change the efficacy of either antibiotic. When used at 10×MIC, there is little effect of phage pre-treatment on antibiotic efficacy. When there is little (CIP, LIN) or no (TET) overlap between confidence intervals, phage pre-treatment actually reduced the antibiotic efficacy (shown in orange ovals on Figure 4). In short, when it had a significant effect, phage pre-treatment improved the efficacy of low concentrations of antibiotics, but it decreased the efficacy of high concentrations of antibiotics. This result is particularly promising for antibiotics prone to toxic side effects, where keeping the concentration low is desirable.

We also addressed the possibility that combining phage with antibiotics would reduce the appearance of antibiotic resistance. In clinical settings, RIF is not used alone in treating infections as resistance commonly ascends during the course of treatment, leading to treatment failure. Indeed, we observed that about half of biofilms treated with RIF alone became turbid, the in vitro analog of treatment failure. To prevent treatment failure due to the ascent of resistance to RIF, RIF is used in combination with other antibiotics to treat infections with *S. aureus* (see (50) for a review). Using phage and RIF, either at the same time or phage first, RIF second, did not increase the efficacy of the antibiotic, but it completely prevented the ascent of RIF resistant bacteria. This observation indicates that one could develop treatment regimens that incorporate phage with RIF in order to reduce the ascent of resistance, while simultaneously diminishing concern of antagonistic interactions between therapeutic agents.

Our results provide evidence supporting the combined use of phage and antibiotics against *S. aureus* biofilms. We observed that the combination of phage and antibiotics prevented the ascent of antibiotic resistant bacteria, and, especially when phage was used first and antibiotic second, it restored efficacy to low concentration antibiotics that would otherwise have been ineffective. These results are promising for the topical use of combinations of antibiotics and phage to treat surface infections, as such surface infections occur in the form of biofilms.

However, more experiments are needed. Future work should examine how general these results are for different strains of *S. aureus*, particularly as different clinical isolates are known to differ in their biofilm-forming abilities. Furthermore, the work could also be expanded to examine the effectiveness of other phages, which may have different pharmacodynamic properties from the single phage tested here. And while promising, these experiments must eventually be conducted in vivo, such as in mouse cutaneous wounds, as the complex dynamics of drug and phage bioavailability, host immune system, bacterial dynamics and kinetics can result in vastly different results from these in vitro experiments. Effectiveness of combined therapy would have to be tested in living model systems in order to pave the way for human clinical trials. Nevertheless, our results are a first step toward a better understanding of the dynamics of treating bacteria with phage and antibiotics.

## Acknowledgements

We thank A. Hanes for obtaining and donating the PYO phage cocktail, Robert Petit III for analyzing the sequencing data, Waqas Chaudhry for his expertise and consulting, Ingrid McCall for making the lab run smoothly, and Bruce Levin, kibbitzer-in-chief, for providing funding for and feedback on this study.

## References

1. Twart F. An investigation on the nature of ultra-microscopic viruses. Lancet. 1915;186(4814):1241–3.

2. d’Herelle F. Sur un microbe invisible antagoniste des bacilles dysentériques (On an invisible microbe antagonistic to dysentery bacilli). Comptes Rendus de l’Académie des Sciences. 1917;165:373–5.

3. d’Herelle F. The Bacteriophage and Its Belhavior. Baltimore: Williams and Wilkins; 1926. 679 p.

4. O’Neill J. Tackling drug-resistant infections globally: final report and recommendation. London; 2016.

5. de Kraker ME, Stewardson AJ, Harbarth S. Will 10 Million People Die a Year due to Antimicrobial Resistance by 2050? PLoS Med. 2016;13(11):e1002184.

6. Abedon ST, Kuhl SJ, Blasdel BG, Kutter EM. Phage treatment of human infections. Bacteriophage. 2011;1(2):66–85.

7. Gorski A, Miedzybrodzki R, Weber-Dabrowska B, Fortuna W, Letkiewicz S, Rogoz P, et al. Phage Therapy: Combating Infections with Potential for Evolving from Merely a Treatment for Complications to Targeting Diseases. Frontiers in microbiology. 2016;7:1515.

8. Henein A. What are the limitations on the wider therapeutic use of phage? Bacteriophage. 2013;3(2):e24872.

9. Shen Y, Barros M, Vennemann T, Gallagher DT, Yin Y, Linden SB, et al. A bacteriophage endolysin that eliminates intracellular streptococci. Elife. 2016;5.

10. Thandar M, Lood R, Winer BY, Deutsch DR, Euler CW, Fischetti VA. Novel Engineered Peptides of a Phage Lysin as Effective Antimicrobials against Multidrug-Resistant Acinetobacter baumannii. Antimicrobial agents and chemotherapy. 2016;60(5):2671–9.

11. Hawkins C, Harper D, Burch D, Anggard E, Soothill J. Topical treatment of Pseudomonas aeruginosa otitis of dogs with a bacteriophage mixture: a before/after clinical trial. Veterinary microbiology. 2010;146(3-4):309–13.

12. Smith HW, Hugggins MB. Successful treatment of experimental Escherichia coli infections in mice using phage: its general superiority over antibiotics. J General Microbiology. 1982;128:307–18.

13. Bull JJ, Levin BR, DeRouin T, Walker N, Bloch CA. Dynamics of success and failure in phage and antibiotic therapy in experimental infections. BMC microbiology. 2002;2:35.

14. Biswas B, Adhya S, Washart P, Paul B, Trostel AN, Powell B, et al. Bacteriophage therapy rescues mice bacteremic from a clinical isolate of vancomycin-resistant Enterococcus faecium. Infect Immun. 2002;70(1):204–10.

15. Ferreira FA, Souza RR, Bonelli RR, Americo MA, Fracalanzza SE, Figueiredo AM. Comparison of in vitro and in vivo systems to study ica-independent Staphylococcus aureus biofilms. Journal of microbiological methods. 2012;88(3):393–8.

16. Alemayehu D, Casey PG, McAuliffe O, Guinane CM, Martin JG, Shanahan F, et al. Bacteriophages phiMR299-2 and phiNH-4 can eliminate Pseudomonas aeruginosa in the murine lung and on cystic fibrosis lung airway cells. mBio. 2012;3(2):e00029–12.

17. Galtier M, De Sordi L, Maura D, Arachchi H, Volant S, Dillies MA, et al. Bacteriophages to reduce gut carriage of antibiotic resistant uropathogens with low impact on microbiota composition. Environmental microbiology. 2016;18(7):2237–45.

18. Roach DR, Leung CY, Henry M, Morello E, Singh D, Di Santo JP, et al. Synergy between the Host Immune System and Bacteriophage Is Essential for Successful Phage Therapy against an Acute Respiratory Pathogen. Cell host & microbe. 2017;22(1):38–47.e4.

19. Waters EM, Neill DR, Kaman B, Sahota JS, Clokie MRJ, Winstanley C, et al. Phage therapy is highly effective against chronic lung infections with Pseudomonas aeruginosa. Thorax. 2017;72(7):666–7.

20. Lang G, Kehr P, Mathevon H, Clavert JM, Sejourne P, Pointu J. Bacteriophage therapy of septic complications of orthopaedic surgery. Revue de chirurgie orthopedique et reparatrice de l’appareil moteur. 1979;65(1):33–7.

21. Schooley RT, Biswas B, Gill JJ, Hernandez-Morales A, Lancaster J, Lessor L, et al. Development and Use of Personalized Bacteriophage-Based Therapeutic Cocktails To Treat a Patient with a Disseminated Resistant Acinetobacter baumannii Infection. Antimicrobial agents and chemotherapy. 2017;61(10).

22. Jennes S, Merabishvili M, Soentjens P, Pang KW, Rose T, Keersebilck E, et al. Use of bacteriophages in the treatment of colistin-only-sensitive Pseudomonas aeruginosa septicaemia in a patient with acute kidney injury-a case report. Critical care (London, England). 2017;21(1):129.

23. Chan BK, Sistrom M, Wertz JE, Kortright KE, Narayan D, Turner PE. Phage selection restores antibiotic sensitivity in MDR Pseudomonas aeruginosa. Scientific reports. 2016;6:26717.

24. Rose T, Verbeken G, Vos DD, Merabishvili M, Vaneechoutte M, Lavigne R, et al. Experimental phage therapy of burn wound infection: difficult first steps. International journal of burns and trauma. 2014;4(2):66–73.

25. Pirnay JP, De Vos D, Verbeken G, Merabishvili M, Chanishvili N, Vaneechoutte M, et al. The phage therapy paradigm: pret-a-porter or sur-mesure? Pharmaceutical research. 2011;28(4):934–7.

26. Debarbieux L, Pirnay JP, Verbeken G, De Vos D, Merabishvili M, Huys I, et al. A bacteriophage journey at the European Medicines Agency. FEMS microbiology letters. 2016;363(2):fnv225.

27. Knouf EG, Ward WE, et al. Treatment of typhoid fever with type specific bacteriophage. J Am Med Assoc. 1946;132:134–8.

28. Desranleau JM. The treatment of typhoid fever by the use of Vi antityphoid bacteriophages; a preliminary report. Can J Public Health. 1948;39(8):317–9.

29. Davies D. Understanding biofilm resistance to antibacterial agents. Nat Rev Drug Discov. 2003;2(2):114–22.

30. Kirby AE, Garner K, Levin BR. The relative contributions of physical structure and cell density to the antibiotic susceptibility of bacteria in biofilms. Antimicrobial agents and chemotherapy. 2012;56(6):2967–75.

31. Hall-Stoodley L, Costerton JW, Stoodley P. Bacterial biofilms: from the Natural environment to infectious diseases. Nat Rev Micro. 2004;2(2):95–108.

32. Stewart PS, Franklin MJ. Physiological heterogeneity in biofilms. Nature reviews Microbiology. 2008;6(3):199–210.

33. Harper DR, Parracho HMRT, Walker J, Sharp R, Hughes G, Werthén M, et al. Bacteriophages and Biofilms. Antibiotics. 2014;3(3):270–84.

34. Danis-Wlodarczyk K, Vandenheuvel D, Jang HB, Briers Y, Olszak T, Arabski M, et al. A proposed integrated approach for the preclinical evaluation of phage therapy in Pseudomonas infections. Scientific reports. 2016;6:28115.

35. Torres-Barceló C, Arias-Sánchez FI, Vasse M, Ramsayer J, Kaltz O, Hochberg ME. A Window of Opportunity to Control the Bacterial Pathogen Pseudomonas aeruginosa Combining Antibiotics and Phages. PLoS one. 2014;9(9):e106628.

36. Chaudhry WN, Concepcion-Acevedo J, Park T, Andleeb S, Bull JJ, Levin BR. Synergy and Order Effects of Antibiotics and Phages in Killing Pseudomonas aeruginosa Biofilms. PLoS one. 2017;12(1):e0168615.

37. Kumaran D, Taha M, Yi Q, Ramirez-Arcos S, Diallo JS, Carli A, et al. Does Treatment Order MatteŘInvestigating the Ability of Bacteriophage to Augment Antibiotic Activity against Staphylococcus aureus Biofilms. Frontiers in microbiology. 2018;9:127.

38. Kwan T, Liu J, DuBow M, Gros P, Pelletier J. The complete genomes and proteomes of 27 Staphylococcus aureus bacteriophages. Proceedings of the National Academy of Sciences of the United States of America. 2005;102(14):5174–9.

39. Jorgensen JH, Ferraro MJ. Antimicrobial susceptibility testing: a review of general principles and contemporary practices. Clinical infectious diseases: an official publication of the Infectious Diseases Society of America. 2009;49(11):1749–55.

40. Johnson PJ, Levin BR. Pharmacodynamics, population dynamics, and the evolution of persistence in Staphylococcus aureus. PLoS Genet. 2013;9(1):e1003123.

41. Team R. RStudio: Integrated Development Environment for R. Boston, MA: RStudio, Inc.; 2015.

42. Bhattacharya M, Wozniak DJ, Stoodley P, Hall-Stoodley L. Prevention and treatment of Staphylococcus aureus biofilms. Expert review of anti-infective therapy. 2015;13(12):1499–516.

43. Mah TF, O’Toole GA. Mechanisms of biofilm resistance to antimicrobial agents. Trends Microbiol. 2001;9(1):34–9.

44. Colvin KM, Gordon VD, Murakami K, Borlee BR, Wozniak DJ, Wong GC, et al. The pel polysaccharide can serve a structural and protective role in the biofilm matrix of Pseudomonas aeruginosa. PLoS pathogens. 2011;7(1):e1001264.

45. Khan W, Bernier SP, Kuchma SL, Hammond JH, Hasan F, O’Toole GA. Aminoglycoside resistance of Pseudomonas aeruginosa biofilms modulated by extracellular polysaccharide. International microbiology: the official journal of the Spanish Society for Microbiology. 2010;13(4):207–12.

46. Abedon ST. Ecology of Anti-Biofilm Agents I: Antibiotics versus Bacteriophages. Pharmaceuticals. 2015;8(3):525–58.

47. Verma V, Harjai K, Chhibber S. Structural changes induced by a lytic bacteriophage make ciprofloxacin effective against older biofilm of Klebsiella pneumoniae. Biofouling. 2010;26(6):729–37.

48. Kim S, Rahman M, Seol SY, Yoon SS, Kim J. Pseudomonas aeruginosa bacteriophage PA1O requires type IV pili for infection and shows broad bactericidal and biofilm removal activities. Applied and environmental microbiology. 2012;78(17):6380–5.

49. Rahman M, Kim S, Kim SM, Seol SY, Kim J. Characterization of induced Staphylococcus aureus bacteriophage SAP-26 and its anti-biofilm activity with rifampicin. Biofouling. 2011;27(10):1087–93.

50. Forrest GN, Tamura K. Rifampin combination therapy for nonmycobacterial infections. Clin Microbiol Rev. 2010;23(1):14–34.

